## Supplementary Material for "Predicting individual traits from models of brain dynamics accurately and reliably using the Fisher kernel"

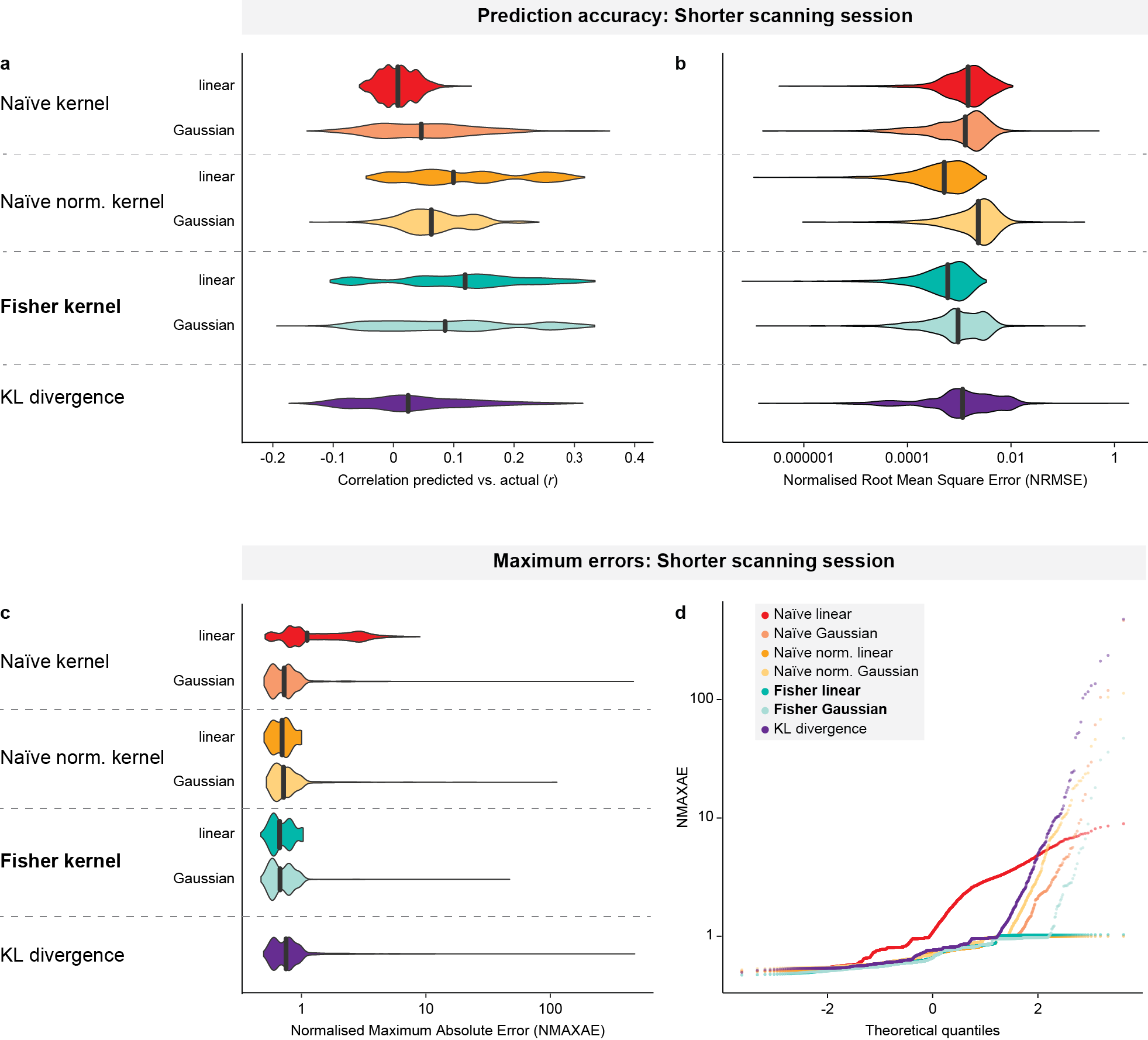

**Figure 1.** Results from using only one scanning session per participant

**Table 1** Behavioural variables

| **Variable no.** | **Column Header** | **Full Display name** | **Assessment** | **HCP Var. no.** |
| --- | --- | --- | --- | --- |
| 1 | Age_in_Yrs | Age in Years | Demographic | 4 |
| 2 | MMSE Score | Mini Mental Status Exam Total Score | Cognitive Status (Mini Mental Status Exam) | 196 |
| 3 | PicSeq_Unadj | NIH Toolbox Picture Sequence Memory Test: Unadjusted Scale Score | Episodic Memory | 222 |
| 4 | PicSeq_AgeAdj | NIH Toolbox Picture Sequence Memory Test: Age-Adjusted Scale Score | Episodic Memory | 223 |
| 5 | CardSort_Unadj | NIH Toolbox Dimensional Change Card Sort Test: Unadjusted Scale Score | Executive Function/ Cognitive Flexibility | 224 |
| 6 | CardSort_AgeAdj | NIH Toolbox Dimensional Change Card Sort Test: Age-Adjusted Scale Score | Executive Function/ Cognitive Flexibility | 225 |
| 7 | Flanker_Unadj | NIH Toolbox Flanker Inhibitory Control and Attention Test: Unadjusted Scale Score | Executive Function/ Inhibition | 226 |
| 8 | Flanker_AgeAdj | NIH Toolbox Flanker Inhibitory Control and Attention Test: Age-Adjusted Scale Score | Executive Function/ Inhibition | 227 |
| 9 | PMAT24_A_CR | Penn Progressive Matrices: Number of Correct Responses (PMAT24_A_CR) | Fluid Intelligence | 228 |
| 10 | PMAT24_A_SI | Penn Progressive Matrices: Total Skipped Items (PMAT24_A_SI) | Fluid Intelligence | 229 |
| 11 | PMAT_A_RTCR | Penn Progressive Matrices: Median Reaction Time for Correct Responses (PMAT24_A_RTCR) | Fluid Intelligence | 230 |
| 12 | ReadEng_Unadj | NIH Toolbox Oral Reading Recognition Test: Unadjusted Scale Score | Language/Reading | 231 |
| 13 | ReadEng_AgeAdj | NIH Toolbox Oral Reading Recognition Test: Age-Adjusted Scale Score | Language/Reading | 232 |
| 14 | PicVocab_Unadj | NIH Toolbox Picture Vocabulary Test: Unadjusted Scale Score | Language/Vocabulary | 233 |
| 15 | PicVocab_AgeAdj | NIH Toolbox Picture Vocabulary Test: Age-Adjusted Scale Score | Language/Vocabulary | 234 |
| 16 | ProcSpeed_Unadj | NIH Toolbox Pattern Comparison Processing Speed Test: Unadjusted Scale Score | Processing Speed | 235 |
| 17 | ProcSpeed_AgeAdj | NIH Toolbox Pattern Comparison Processing Speed Test: Age-Adjusted Scale Score | Processing Speed | 236 |
| 18 | VSPLOT_TC | Variable Short Penn Line Orientation: Total Number Correct (VSPLOT_TC) | Spatial Orientation | 251 |
| 19 | VSPLOT_CRTE | Variable Short Penn Line Orientation: Median Reaction Time Divided by Expected Number of Clicks for Correct (VSPLOT_CRTE) | Spatial Orientation | 252 |
| 20 | VSPLOT_OFF | Variable Short Penn Line Orientation: Total Positions Off for All Trials (VSPLOT_OFF) | Spatial Orientation | 253 |
| 21 | SCPT_TP | Short Penn Continuous Performance Test: True Positives = Sum of CPN_TP and CPL_TP (SCPT_TP) | Sustained Attention | 254 |
| 22 | SCPT_TN | Short Penn Continuous Performance Test: True Negatives = Sum of CPN_TN and CPL_TPN (SCPT_TN) | Sustained Attention | 255 |
| 23 | SCPT_FP | Short Penn Continuous Performance Test: False Positives = Sum of CPN_FP and CPL_FP (SCPT_FP) | Sustained Attention | 256 |
| 24 | SCPT_FN | Short Penn Continuous Performance Test: False Negatives = Sum of CPN_FN and CPL_FN (SCPT_FN) | Sustained Attention | 257 |
| 25 | SCPT_TRPRT | Short Penn Continuous Performance Test: Median Response Time for True Positive Responses (SCPT_TPRT) | Sustained Attention | 258 |
| 26 | SCPT_SEN | Short Penn Continuous Performance Test: Sensitivity = SCPT_TP/(SCPT_TP + SCPT_FN) (SCPT_SEN) | Sustained Attention | 259 |
| 27 | SCPT_SPEC | Short Penn Continuous Performance Test: Specificity = SCPT_TN/(SCPT_TN + SCPT_FP) (SCPT_SPEC) | Sustained Attention | 260 |
| 28 | SCPT_LRNR | Short Penn Continuous Performance Test: Longest Run of Non-Responses (SCPT_LRNR) | Sustained Attention | 261 |
| 29 | IWRD_TOT | Penn Word Memory Test: Total Number of Correct Responses (IWRD_TOT) | Verbal Episodic Memory | 262 |
| 30 | IWRD_RTC | Penn Word Memory Test: Median Reaction Time for Correct Responses (IWRD_RTC) | Verbal Episodic Memory | 263 |
| 31 | ListSort_Unadj | NIH Toolbox List Sorting Working Memory Test: Unadjusted Scale Score | Working Memory | 264 |
| 32 | ListSort_AgeAdj | NIH Toolbox List Sorting Working Memory Test: Age-Adjusted Scale Score | Working Memory | 265 |
| 33 | Language_Task_Acc | Language Task OVERALL Accuracy | Language Task | 510 |
| 34 | Relational_Task_Acc | Relational Task OVERALL Accuracy | Relational Task | 518 |
| 35 | WM_Task_Acc | Working Memory Task OVERALL Accuracy | Working Memory Task | 545 |

**Table 2** Summary of model performance. The table shows the average performance and the range of the correlation between model-predicted and actual values (r) in deconfounded space, the coefficient of determination (R^2^) in deconfounded space, and of the normalised maximum errors (NMAXAE) in original space.

| **Method** | | **Correlation coefficient (*r*)** | | | | **Coefficient of determination (*R^2^*)** | | | | **NMAXAE** | | | |
| --- | --- | --- | --- | --- | --- | --- | --- | --- | --- | --- | --- | --- | --- |
|  | | **min.** | **mean** | **median** | **max.** | **min.** | **mean** | **median** | **max.** | **min.** | **mean** | **median** | **max.** |
| **Fisher** | Linear | -0.28 | 0.19 | 0.19 | 0.67 | -4.89 | -0.01 | 0.01 | 0.40 | 0.04 | 0.50 | 0.48 | 1.00 |
|  | Gaussian | -0.50 | 0.17 | 0.16 | 0.68 | -9.99E+02 | -0.01 | 0.02 | 0.41 | 0.02 | 0.49 | 0.48 | 18.28 |
| **Naïve** | Linear | -0.41 | 0.05 | 0.05 | 0.52 | -16.04 | -0.05 | 0.00 | 0.26 | 0.02 | 0.55 | 0.50 | 10.27 |
|  | Gaussian | -0.34 | 0.11 | 0.10 | 0.63 | -5.59E+03 | -0.19 | 0.00 | 0.38 | 0.03 | 0.51 | 0.49 | 42.81 |
| **Naïve norm.** | Linear | -0.28 | 0.15 | 0.14 | 0.71 | -2.06 | 0.00 | 0.01 | 0.40 | 0.03 | 0.50 | 0.48 | 1.00 |
|  | Gaussian | -0.36 | 0.09 | 0.09 | 0.62 | -2.37E+08 | -6.85E+03 | 0.00 | 0.39 | 0.02 | 0.96 | 0.49 | 1.22E+04 |
| **KL div.** | | -0.32 | 0.16 | 0.16 | 0.65 | -1.15E+05 | -6.30 | 0.01 | 0.41 | 0.02 | 0.54 | 0.49 | 1.03E+02 |
| **ta KL div.** | | -0.31 | 0.19 | 0.19 | 0.67 | -4.21E+04 | -2.27 | 0.03 | 0.45 | 0.02 | 0.53 | 0.49 | 98.65 |
| **Log-Euclidean** | | -0.48 | 0.14 | 0.13 | 0.61 | -0.24 | 0.03 | 0.01 | 0.31 | 0.02 | 0.50 | 0.48 | 0.99 |
| **Ridge reg.** | | -0.41 | 0.11 | 0.10 | 0.66 | -9.30 | 0.02 | 0.00 | 0.33 | 0.02 | 0.51 | 0.48 | 5.89 |
| **Ridge reg. (Riem.)** | | -0.31 | 0.22 | 0.23 | 0.74 | -0.20 | 0.07 | 0.04 | 0.49 | 0.02 | 0.49 | 0.47 | 0.99 |
| **Selected Edges** | | -0.36 | 0.08 | 0.08 | 0.52 | -2.96 | -0.02 | -0.01 | 0.21 | 0.02 | 0.50 | 0.49 | 0.99 |

­

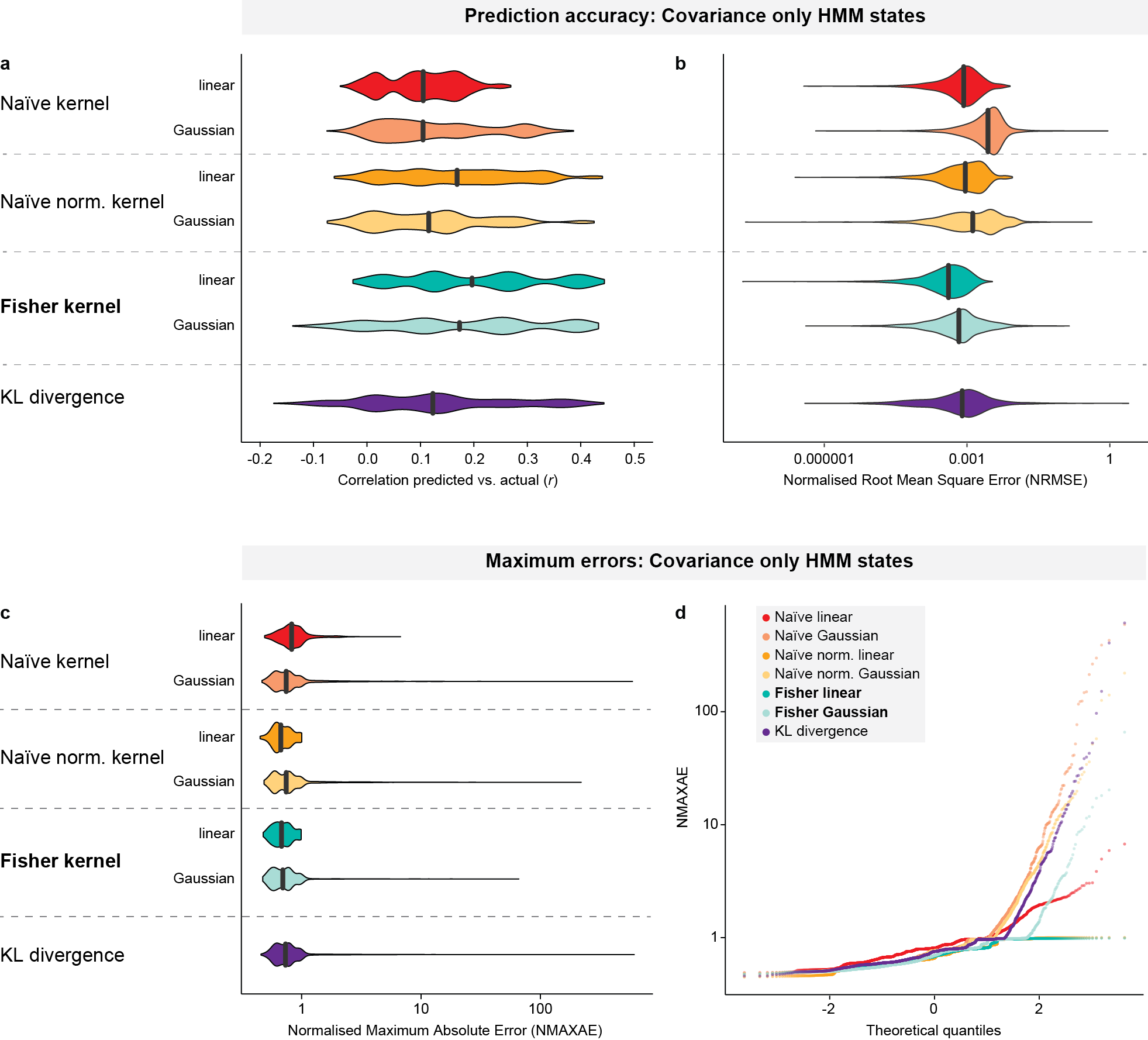

**Figure 2** Results from HMM where states are defined only in terms of covariance and mean is set to 0.

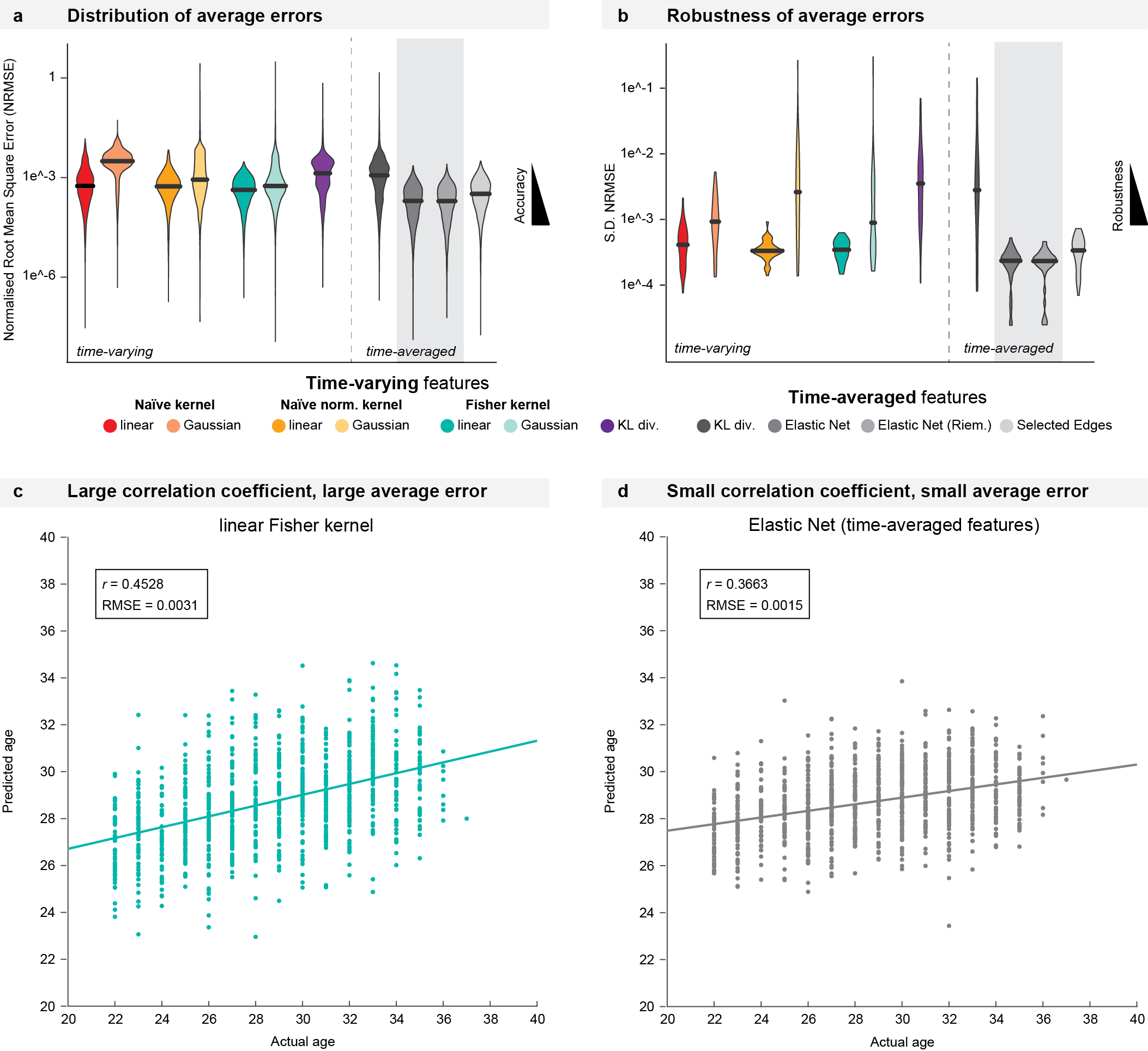

**Figure 3** Average errors. a), b) Distribution and robustness of average errors. c), d) While a larger correlation coefficient indicates a stronger relationship between the predictor and the predicted variable (c), a smaller average error (d) can be achieved by predicting closely around the mean. This is the case for the Elastic Net models and the Selected Edges model.

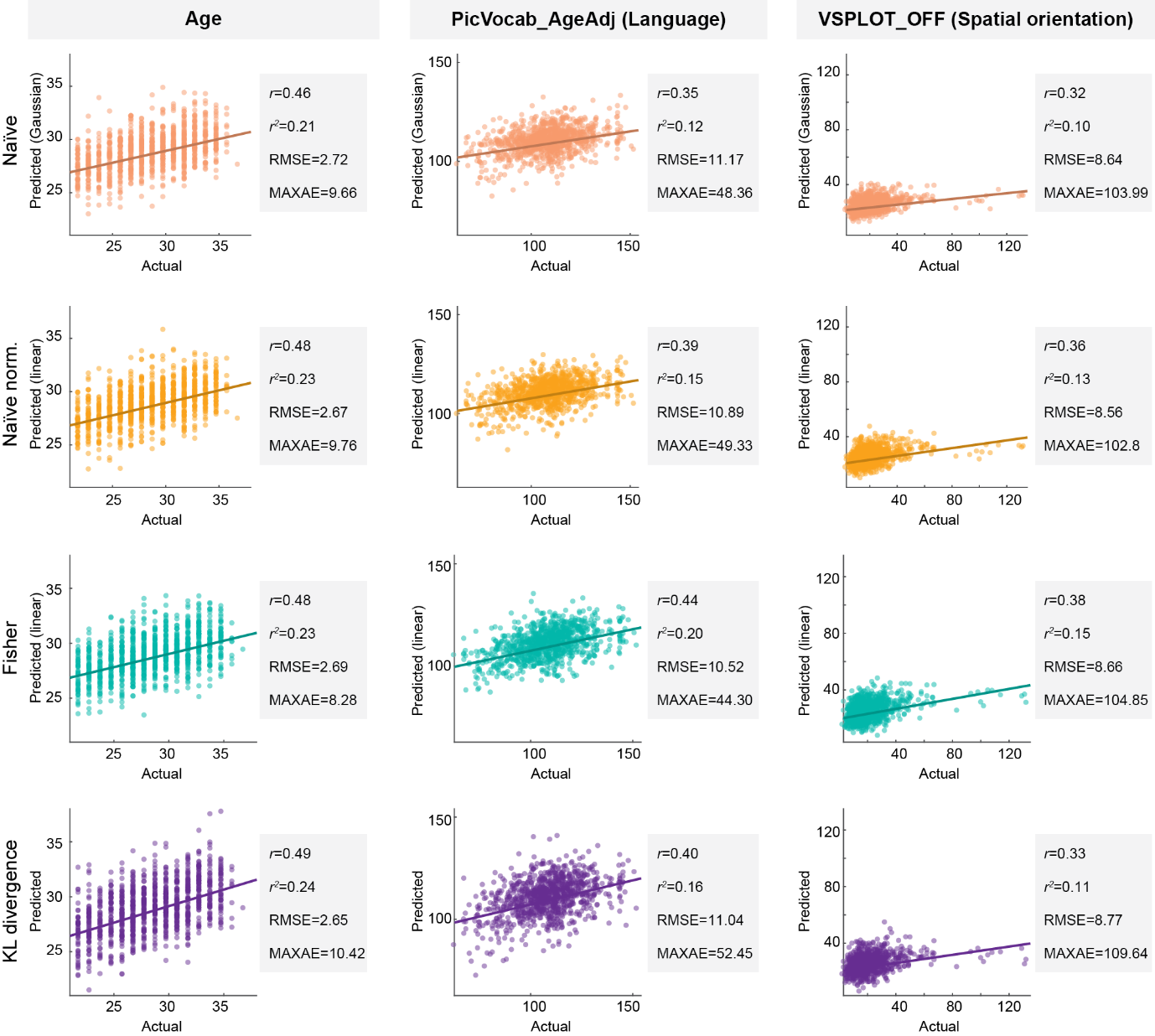

**Figure 4** Best predictions for three example variables from each kernel. The plot shows a single run from each kernel that produced the highest prediction accuracy for the three example variables. In the single best runs of these variables, all kernels produce similarly accurate results. Note however, that, since the naïve kernel and the KL divergence model were less robust, they only produced fairly accurate predictions as shown here in few selected runs, while they produced large errors in other runs. The RMSE and maximum error values reported beside the plots are here not normalised (i.e. the errors are on the same scale as the variable).

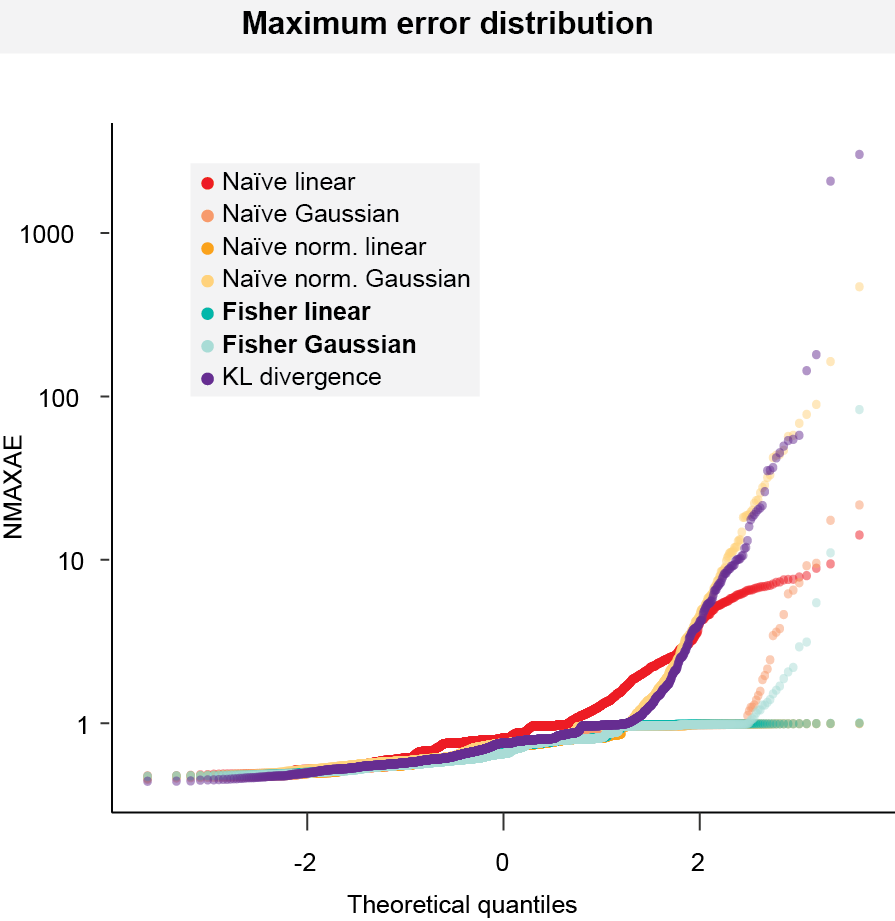

**Figure 5** Quantile-quantile plot of normalised maximum errors. The plot highlights how the differences between the kernels lie mainly in the tails of the distribution, where the KL divergence and the Gaussian naïve normalised kernel have the longest tails.

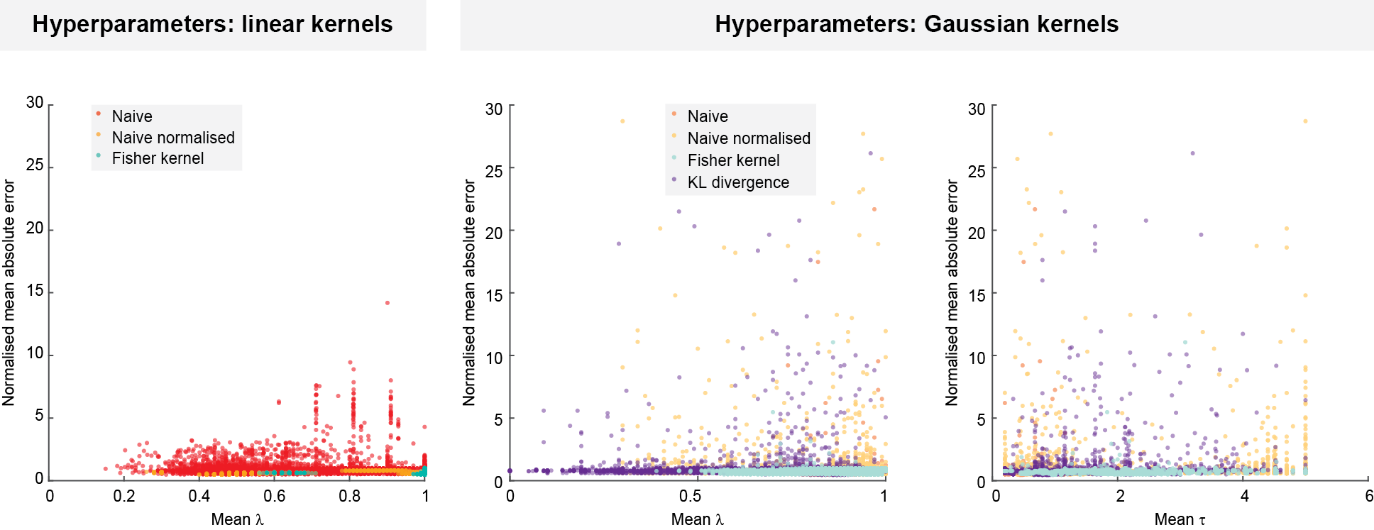

**Figure 6** Effect of hyperparameters on model errors
